# Primary CD34^+^ cells of patients with VEXAS syndrome are highly sensitive to targeted treatment with TAK-243 and pevonedistat

**DOI:** 10.1101/2025.05.22.653244

**Authors:** Daniel Nowak, Marie Demmerle, Alessa Klär, Alexander Streuer, Vladimir Riabov, Eva Altrock, Felicitas Rapp, Ann-Christin Belzer, Teresa Klink, Meret Hahn, Franziska Hofmann, Verena Nowak, Nadine Weimer, Julia Obländer, Iris Palme, Melda Göl, Ali Darwich, Laurenz Steiner, Mohammed Abba, Georgia Metzgeroth, Wolf-Karsten Hofmann, Nanni Schmitt

## Abstract

Vacuoles, E1 enzyme, X-linked, Autoinflammatory, Somatic (VEXAS) syndrome is an autoinflammatory disease characterized by somatically acquired *UBA1* mutations in hematopoietic stem cells. VEXAS patients present with significant therapeutic challenges such as severe treatment-resistant inflammation and predisposition to myeloid malignancies. In this study, we evaluated the efficacy of TAK-243, a UBA1 inhibitor, and pevonedistat, a NEDD8 inhibitor, *in vitro* on primary CD34^+^ hematopoietic stem and progenitor cells (HSPCs) from patients with VEXAS syndrome (n=5), myelodysplastic neoplasms (MDS) (n=10), as well as healthy controls (n=10). Our findings revealed that VEXAS patient-derived HSPCs exhibited significantly higher sensitivity to both TAK-243 (IC50: 22 nM) and pevonedistat (IC50: 553 nM) compared to MDS (IC50: 133 nM and 1048 nM, respectively) and healthy controls (IC50: 192 nM and 1107 nM, respectively) as demonstrated by reduced cell viability. Furthermore, VEXAS CD34^+^ cells exhibited significantly higher levels of apoptosis after treatment with both inhibitors. These results suggest that TAK-243 and pevonedistat selectively target *UBA1*-mutated progenitor cells, and indicate a broad therapeutic window in which malignant cells are eradicated but healthy hematopoiesis is not yet compromised. This study provides a strong rationale for further clinical investigation of TAK-243 and pevonedistat as targeted therapies for VEXAS patients, offering hope for improved management of this clinically challenging disorder and contributing to our understanding of its underlying molecular mechanisms.

## Introduction

Vacuoles, E1 enzyme, X-linked, Autoinflammatory, Somatic (VEXAS) syndrome is a systemic autoinflammatory disease characterized by somatically acquired inactivating mutations of the *UBA1* gene in hematopoietic stem and progenitor cells.^1^ *UBA1* encodes the Ubiquitin Like Modifier Activating Enzyme 1, which plays a crucial role in the post-translational modification processes of ubiquitination and atypical neddylation responsible for regulation of protein degradation and cellular homeostasis.^2,3^ VEXAS syndrome predominantly affects men in late adulthood, between 66-76 years of age, and is characterized by severe, treatment-refractory inflammation, including recurrent fever, chondritis, neutrophilic dermatosis, alveolitis, vasculitis and cytopenias. Additionally, VEXAS patients have a predisposition for the development of myeloid malignancies such as myelodysplastic neoplasms (MDS).^4^ Morphologically, VEXAS presents with hypercellular bone marrow, varying degrees of dysplasia, macrocytic anemia and, most prominently, vacuolated myeloid and erythroid precursors.^5^

Given the recent discovery of VEXAS syndrome, standardized treatment options are not yet established, leaving patients with high morbidity and mortality.^6^ Disease management primarily involves anti-inflammatory therapies such as high-dose corticosteroids, which are associated with major toxicity.^7^ For patients with concomitant myeloid neoplasia, treatment with the hypomethylating agent azacytidine and JAK inhibitors achieved mixed results, while high incidences of severe infections were found to be frequent adverse events.^8-10^ The only potentially curative approach is an allogeneic stem cell transplantation.

Complete loss of UBA1 function is non-viable to cells and organisms.^11^ The most common mutations in VEXAS syndrome are *UBA1 p*.*M41T, UBA1 p*.*M41V* and *UBA1 p*.*M41L*, which affect methionine 41 of exon 3 and lead to isoforms of UBA1 with considerably reduced catalytic activity.^12^, However, current scientific data suggest that residual catalytic activity is retained in cells of VEXAS, which enables affected myeloid progenitors to survive and gain a selective clonal advantage.^1^ Therefore, a particularly promising approach for molecularly targeted treatment of VEXAS syndrome involves drugs targeting the ubiquitin proteasome system. Two potential candidates are the inhibitors TAK-243 (MLN7243), which targets E1 ubiquitin activating enzymes (UAEs), and pevonedistat (TAK-924, MLN4924), which targets NEDD8 activating enzymes (NAEs).^13,14^

Recent research by Chiaramida and colleagues demonstrated the therapeutic potential of TAK-243 using a murine Uba1M41L knock-in cell line model that replicated VEXAS syndrome features.^15^ Building on these findings, in this study, we conducted *in vitro* testing of TAK-243 and pevonedistat on primary CD34^+^ cells from patients with VEXAS syndrome (n=5) and MDS (n=10), as well as from healthy individuals (n=10), to explore their clinical efficacy.

## Results

Examination of the clinical characteristics of the five VEXAS patients revealed several common features of VEXAS syndrome (**Supplementary Table S1**). At the time of bone marrow aspiration, the mean age was 72.4 years. All VEXAS patients presented with inflammatory symptoms, elevated C-reactive protein levels and MDS as hematologic comorbidity. Four of five VEXAS patients received prednisolone. Vacuolization of granulopoiesis and/or erythropoiesis was observed in all patients. All five VEXAS patients had missense variants of codon 41: two patients harbored the hot spot mutation p.Met41Leu (variant allele frequency (VAF) 83% and 82%), while the other three carried the p.Met41Thr variant (VAF 90%, 89% and 36%). Further, panel sequencing revealed the VEXAS-typical co-mutation *DNMT3A* in two patients (VAF 40% and 15%). The presence of UBA1 variants in CD34^+^ cells from VEXAS patients was confirmed using droplet digital PCR (**Supplementary Figure S1**).

To assess the therapeutic potential of TAK-243 and pevonedistat, we isolated CD34^+^ cells from bone marrow mononuclear cells of VEXAS and MDS patients, as well as healthy controls. Our findings revealed that primary VEXAS CD34^+^ cells exhibit significantly higher sensitivity to TAK-243, with an IC50 of 22 nM, compared to MDS (133 nM, p=0.0047) and healthy cells (192 nM, p=0.0005) as demonstrated by decreasing cell viability (**Figure 1A+B**). Further, MDS cells were more susceptible to treatment with TAK-243 than healthy CD34+ cells. Pevonedistat also showed increased efficacy in VEXAS cells (IC50 553 nM) compared to MDS (1048 nM, p=0.0004) and healthy CD34^+^ cells (1107 nM, p=0.0002) (**Figure 1C+D**). IC50 results for pevonedistat in MDS and healthy CD34^+^ cells were nearly identical. Overall, MDS and healthy samples only responded to higher concentrations of TAK-243 (>100 nM) and pevonedistat (>500 nM). Notably, minimal variability was observed in the IC50 results for TAK-243 and pevonedistat across all samples of each entity (**Supplementary Figure 2A-F**).

**Figure 1.**
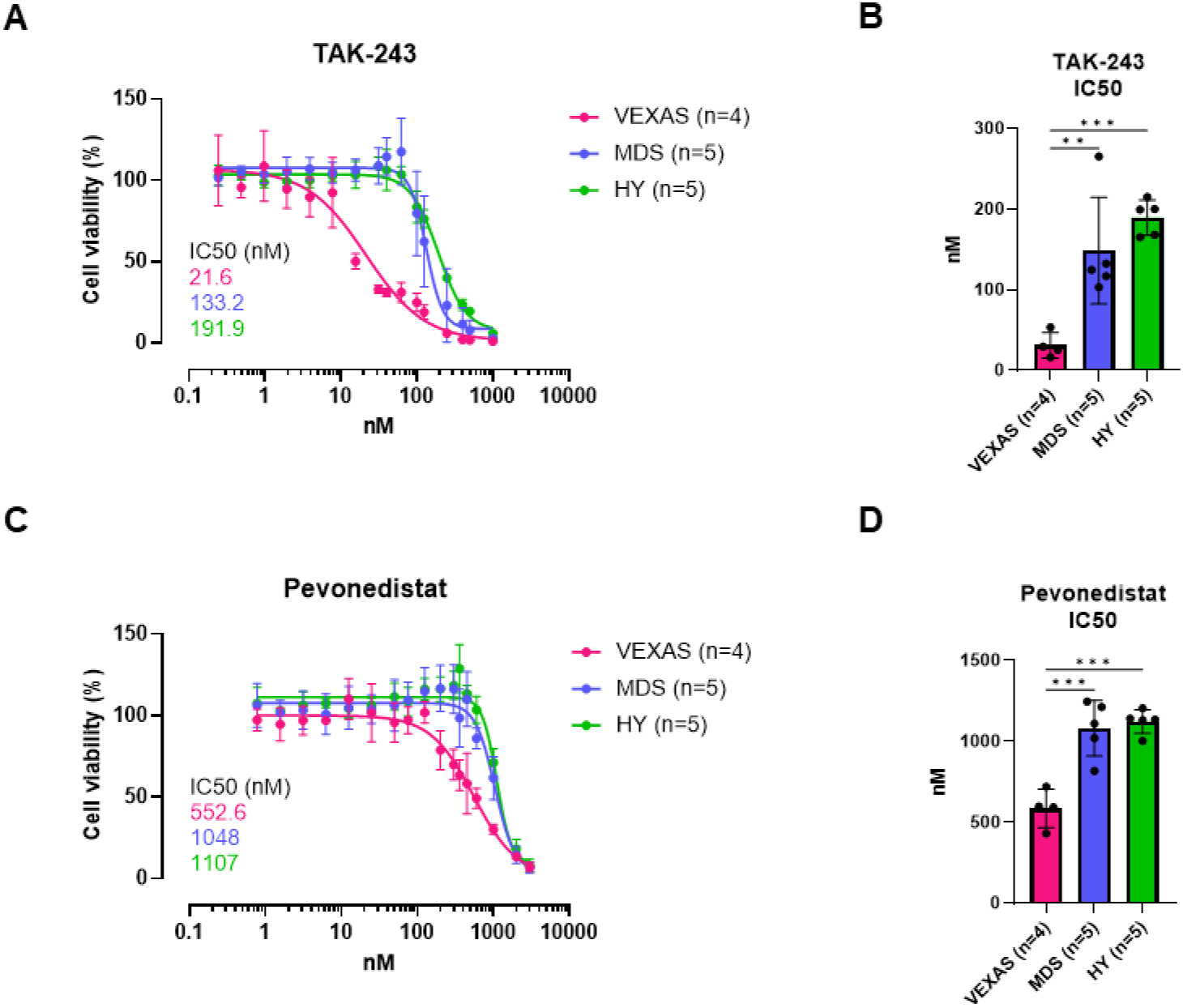
VEXAS CD34^+^ cells are highly sensitive to treatment with TAK-243 and pevonedistat compared to MDS and healthy samples. Dose-response experiments of primary VEXAS (n=4; magenta), MDS (n=5; blue) and healthy (n=5; green) CD34^+^ cells treated with TAK-243 or pevonedistat. Cell viability was measured 48 hours after treatment using CellTiter-Glo® Luminescent Cell Viability Assay. **A**+**C** Dose response curves of TAK-243 (**A**) and pevonedistat (**C**). **B**+**D** IC50s of TAK-243 (**B**) and pevonedistat (**D**). Data were analyzed using nonlinear regression (**A**+**C**) and two-tailed unpaired t-test (**B**+**D**) using Prism 9 (GraphPad Software), and are represented as mean ± SD. **p≤0.01, ***p≤0.001

Interestingly, CD34^+^ cells of all three groups demonstrated significantly higher sensitivity to TAK-243 than to pevonedistat, as evidenced by lower IC50 values for cell viability (p<0.0001 for all) (**Supplementary Figure S3A-F**). This difference can most probably be attributed to the fact that pevonedistat, unlike TAK-243, does not directly interfere with the ubiquitination process, but as an inhibitor of NAE affects neddylation. Although UBA1 is also involved in the process of atypical neddylation, mutations in UBA1 likely have a weaker effect on this specific pathway.

Furthermore, we could demonstrate that VEXAS CD34^+^ cells exhibited significantly higher levels of apoptosis after treatment with TAK-243 and pevonedistat compared to both MDS (TAK-243: IC75 p=0.0007; pevonedistat: IC25 p=0.035, IC50 p=0.0031, IC75 p=0.0049) and healthy CD34^+^ cells (TAK-243: IC50 p=0.0005, IC75 p=0.0004; pevonedistat: IC25 p=0.0064, IC50 p=0.0027, IC75 p=0.0077) (**Figure 2A+B**). The differences between MDS and healthy CD34^+^ cells were again not significant for any of the concentrations. These results provide further evidence that TAK-243 is more effective than pevonedistat in killing VEXAS CD34^+^ cells. Due to the high levels of apoptosis, it can be assumed that TAK-243 and pevonedistat primarily induce apoptotic cell death, which is probably triggered by the disruption of the ubiquitin-proteasome pathway.

**Figure 2.**
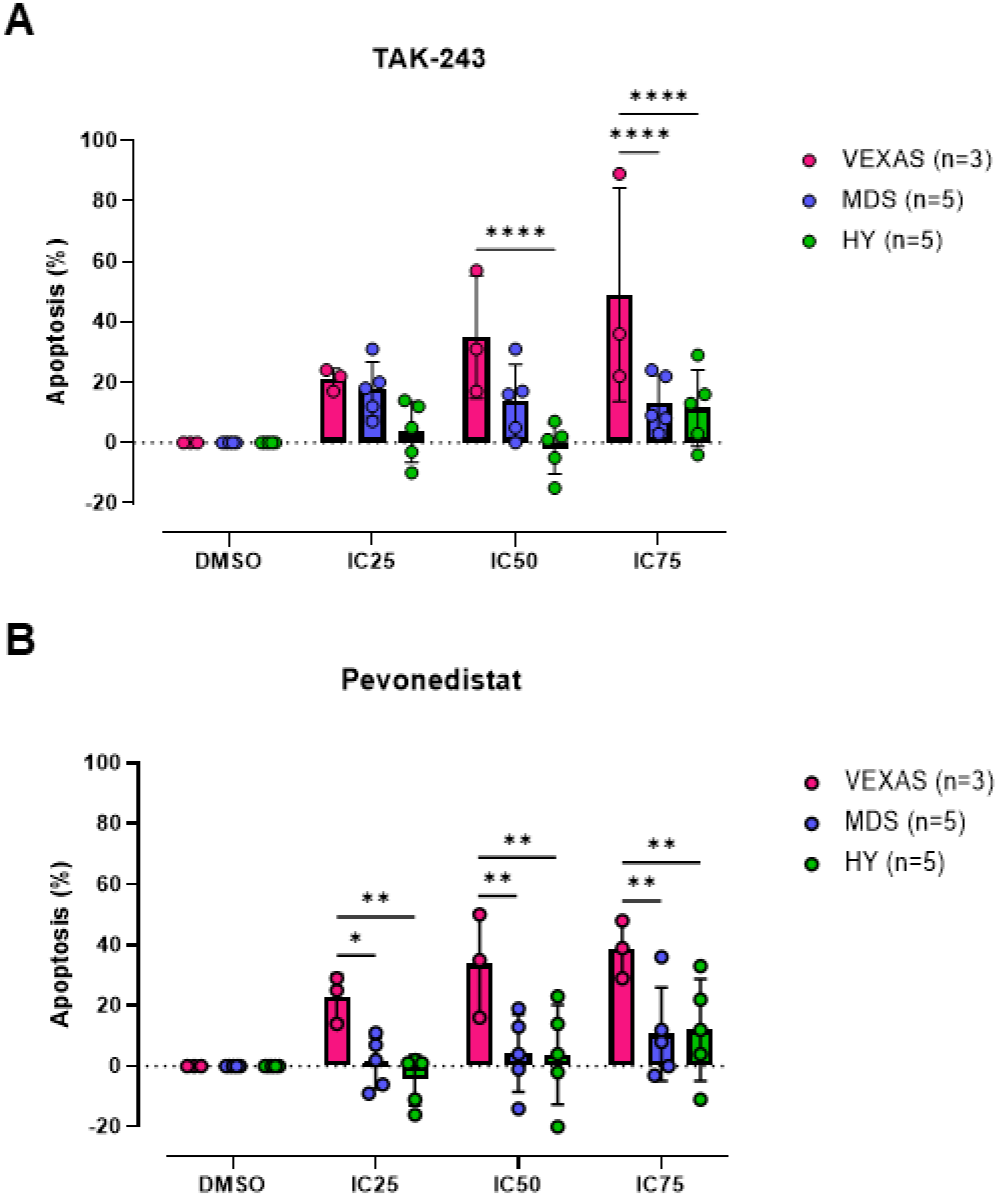
VEXAS CD34^+^ cells exhibit a higher level of apoptosis following treatment with TAK-243 and pevonedistat compared to MDS and healthy samples. Dose-response experiments of primary VEXAS (n=3; magenta), MDS (n=5; blue) and healthy (n=5; green) CD34^+^ cells treated with TAK-243 (**A**) or pevonedistat (**B**). IC25, IC50 and IC75 for CD34^+^ cells of VEXAS patients calculated from **Figure 1** were used for treatment. The percentage of apoptotic cells was measured after 48 hours using Caspase-Glo® 3/7 Assay. Data were analyzed using ordinary two-way ANOVA using Prism 9 (GraphPad Software), and are represented as mean ± SD. *p≤0.05, **p≤0.01, ****p≤0.0001

## Discussion

Our study provides compelling preclinical evidence supporting the potential of TAK-243 and pevonedistat as molecularly targeted therapies for VEXAS syndrome. Primary CD34^+^ cells from VEXAS patients exhibited significantly higher sensitivity to both TAK-243 and pevonedistat compared to cells from MDS patients and healthy controls as demonstrated by lower cell viabilities and higher levels of apoptosis. Particularly in the case of TAK-243, this indicates a broad therapeutic window in which malignant hematopoiesis could be eradicated but healthy hematopoiesis is not yet affected. Further, our results suggest that treatment with these substances selectively targets UBA1-mutated myeloid progenitor cells, which are the origin of the inflammatory disease, by inducing synthetic lethality, while sparing healthy hematopoietic progenitors. This increased selective susceptibility of VEXAS CD34^+^ cells to these inhibitors presents a promising therapeutic approach for a more effective and less toxic treatment for this challenging disorder.

In summary, we here provide a strong rationale for further clinical investigation of TAK-243 and pevonedistat as potential targeted therapies for patients with VEXAS syndrome. These findings offer hope for improving the management of this recently discovered disorder and contribute to our understanding of its underlying molecular mechanisms. Future studies should also explore combination therapies with standard MDS treatments for patients with concomitant myeloid neoplasms.

## Methods

### Primary samples

Primary bone marrow samples from patients with VEXAS (n=5) or MDS (n=10) were obtained from residual diagnostic bone marrow aspirations. Bone marrow of healthy, age-matched controls (n=10) was isolated from femoral heads received after hip replacement surgery. All samples were collected after obtaining the patients’ written informed consent and in accordance with the Declaration of Helsinki. Detailed information on patients and controls are provided in **Supplementary Tables S1+2**. Bone marrow mononuclear cells were isolated by red cell lysis. Enrichment of CD34+ cells from mononuclear cells by magnetic cell separation was performed using the ‘CD34 MicroBead’ kit (Miltenyi Biotech) yielding a purity of at least 90%. Enriched CD34+ cells were viably frozen until further use.

### Droplet digital PCR

DNA from CD34^+^ cells was isolated using the ‘AllPrep DNA/RNA’ kit (QIAGEN). Detection of UBA1 variants by digital droplet PCR was performed using the ‘QX200 Droplet Digital PCR System’ (Bio-Rad Laboratories). Primers (forward: 5’-CTCCACTCCTGTGTGTCT-3’, reverse: 5’-GTAAAGGCCCTCGTCTATGT-3’) and probes for UBA1 wild type (5’-HEX-CTAGGGAATGGC-3’), UBA1 p.Met41Thr (5’-FAM–CTAGGGAACGGC-3’) and UBA1 p.Met41Leu (5’-FAM–CTAGGGACTGGC-3’) were purchased as ready-to-use mix in a ratio of 900:575 nM (Bio-Rad Laboratories). Droplets were generated using the ‘QX200 Droplet Generator’ (Bio-Rad Laboratories). PCR reactions were performed using the ‘C1000 Touch Thermal Cycler with 96–Deep Well Reaction Module’ (Bio-Rad Laboratories). Droplets were counted using the ‘QX200 Droplet Reader’ (Bio-Rad Laboratories). Analysis was performed using the ‘QX Manager’ software (Bio-Rad Laboratories, v2.0.0).

### Cell viability and apoptosis assays

After thawing, CD34+ cells were cultured in StemSpan™ SFEM II (STEMCELL Technologies) plus cytokines (10 ng/ml FGF-1, 50 ng/ml FLT3-L, 50 ng/ml SCF, 10 ng/ml TPO) and 1% pen-strep overnight. To set up the assays, 2,500 CD34+ cells per well in StemSpan™ SFEM II plus cytokines in replicates of 3-5 were treated with different concentrations of TAK-243 (range: 0.24–1,000 nM) and pevonedistat (range: 0.78–3,000 nM) or vehicle (DMSO) for 48 hours. Cell viability and apoptosis were then assessed using the CellTiter-Glo® Luminescent Cell Viability Assay or Caspase-Glo® 3/7 Assay (Promega), respectively. The luminescence was measured using the ‘Tecan Infinite 200 PRO 8’ microplate reader (Tecan Group Ltd.). All results were normalized according to the DMSO results.

### Statistical analysis

Statistical analysis was performed using Prism 9.4.1 (GraphPad Software, San Diego, CA, USA). P values less than 0.05 were considered significant (n.s.: not significant, statistically significant *p<0.05, **p<0.01, ***p<0.001, ****p<0.0001).

## Data availability

Data were generated by the authors and available on request.

## Acknowledgment

This work was supported by the “Forum Gesundheitsstandort Baden-Württemberg”, Projektvorhaben „Identifizierung und Nutzung molekularer und biologischer Muster für die individuelle Krebsbehandlung” BW 4-5400/136/1 (D.N.), the German Cancer Aid Foundation (Deutsche Krebshilfe, 70113953) (D.N.), Medical Faculty Mannheim of the Heidelberg University „SEED” (N.S.), “Torsten Haferlach Leukämiediagnostik Stiftung” (N.S.) and Health + Life Science Alliance Heidelberg Mannheim (V.R.). This work was supported by state funds approved by the State Parliament of Baden-Württemberg.

## Contribution

N.S., D.N. and M.D. designed the study, analyzed and interpreted data, and wrote the manuscript; N.S., M.D. and A.K. performed experiments; A.S. executed bioinformatic analyses; V.R., E.A., F.R., A.-C.B., T.K., M.H., F.H., V.N., N.W., J.O., I.P. and M.G. provided technical assistance; A.D. provided primary material from healthy controls; A.S., M.A. and G.M. provided patient material and clinical data; D.N. and W.-K.H. supervised the whole study and provided research infrastructure.

