## Supplementary Material for "Primary CD34^+^ cells of patients with VEXAS syndrome are highly sensitive to targeted treatment with TAK-243 and pevonedistat"

**Supplementary Table S1. Detailed genetic and clinical characteristics of VEXAS patients.** CRP, C-reactive protein; ESA, erythropoiesis-stimulating agent; Hb, hemoglobin; IPSS-R/IPSS-M, Revised/Molecular International Prognostic Scoring System for MDS; MDS, myelodysplastic neoplasms; MDS-LB, MDS with low blasts (WHO 2022); PLT, platelets; WBC, white blood cells; VAF, variant allele frequency; *at time of bone marrow aspiration; **time from first diagnosis to death/last follow-up; *** 1) viability assay, 2) apoptosis assay.

| Patient ID | Sex | Age* | Overall survival** | Clinical manifestation* | Blood parameters* | Cytology* | Karyotype* | Mutations* | Treatment* | Used for assay*** |
|---|---|---|---|---|---|---|---|---|---|---|
| **VEXAS1** | male | 74 | 14 months | - MDS-LB (IPSS-R: low, IPSS-M: low)<br>- Fever<br>- Papular and pustular rash on whole body<br>- Petechiae on the lower legs | - Anemia (Hb: 9.2 g/dl)<br>- Leukopenia (WBC: 4.17x$10^9$/L)<br>- Elevated CRP (82 mg/l) | - Vacuolization in the immature erythroid precursors<br>- Dysplasia in erythropoiesis and granulopoiesis | 46, XY [10] | - UBA1 p.Met41Thr (VAF 89%)<br>- DNMT3A (VAF 40%) | - Prednisolone | 1, 2 |
| **VEXAS2** | male | 73 | 40 months | - MDS-LB (IPSS-R: low, IPSS-M: low)<br>- Progressive cutaneous leukemic infiltrates (Sweet syndrome) | - Anemia (Hb: 9.3 g/dl)<br>- Thrombocytopenia (PLT: 25x$10^9$/L)<br>- Elevated CRP (45.3 mg/l) | - Vacuolization in the erythroid and myeloid lineage<br>- Dysplasia in erythropoiesis, granulopoiesis and megakaryopoiesis | 46, XY [20] | - UBA1 p.Met41Leu (VAF 82%) | - ESA<br>- Asunercept<br>- Iron chelator | 1 |
| **VEXAS3** | male | 74 | 10+ months | - MDS-LB (IPSS-R: intermediate, IPSS-M: moderate low) | - Anemia (Hb: 6.9 g/dl)<br>- Leukopenia (WBC: 2.83x$10^9$/L)<br>- Thrombocytopenia (PLT: 72x$10^9$/L)<br>- Elevated CRP (61 mg/l) | - Vacuolization in promyelocytes<br>- Dysplasia in megakaryopoiesis | 46,XY [20] | - UBA1 p.Met41Leu (VAF 83%) | - Prednisolone<br>- Azathioprine | 1 |
| **VEXAS4.1 (= VEXAS4.2)** | male | 71 | 15+ months | - MDS-LB (IPSS-R: very low, IPSS-M: very low)<br>- Fever of unknown origin<br>- Arteritic anterior ischemic optic neuropathy<br>- Maculopapular rash on the left upper arm<br>- Mesenteric panniculitis<br>- Recurrent polychondritis | - Anemia (Hb: 12.4 g/dl)<br>- Leukopenia (WBC: 3.41x$10^9$/L)<br>- Elevated CRP (37 mg/l) | - Vacuolization in the precursors of granulopoiesis, predominantly in promyelocytes and proerythroblasts<br>- Dysplasia in erythropoiesis, granulopoiesis and megakaryopoiesis | 46,XY [20] | - UBA1 p.Met41Thr (VAF 90%) | - Prednisolone | 1 |

| | | | | | | | | | | |
|---|---|---|---|---|---|---|---|---|---|---|
| **VEXAS4.2 (= VEXAS4.1)** | male | 72 | 15+ months | (see above) | - Anemia (Hb: 7.4 g/dl)<br>- Elevated CRP (90.7 mg/l) | - Vacuolization in (pro)myelocytes<br>- Dysplasia in granulopoiesis and megakaryopoiesis | 46,XY [20] | - UBA1 p.Met41Thr (VAF 87%) | - Prednisolone<br>- Ruxolitinib | 2 |
| **VEXAS5** | male | 70 | 4+ months | - MDS-LB (IPSS-R: very low, IPSS-M: very low)<br>- Recurrent polychondritis | - Elevated CRP (16.9 mg/l) | - Vacuolization in erythropoiesis and granulopoiesis<br>- Dysplasia in erythropoiesis, granulopoiesis and megakaryopoiesis | 46,XY [20] | - UBA1 p.Met41Thr (VAF 36%)<br>- SMC3 (VAF 16%)<br>- DNMT3A (VAF 15%) | - Prednisolone | 2 |

**Supplementary Table S2. Key clinical data of MDS patients and healthy controls.** ESA, erythropoiesis-stimulating agent; HY, healthy; IPSS-R/IPSS-M, Revised/Molecular International Prognostic Scoring System for MDS; int, intermediate; MDS, myelodysplastic neoplasms; MDS-LB, MDS with low blasts; MDS-SF3B1, MDS with SF3B1 mutation; WHO 2022, WHO classification of MDS 2022; *at time of bone marrow aspiration; ** 1) viability assay, 2) apoptosis assay.

| Patient ID | Sex | Age* | Diagnosis (WHO 2022)* | IPSS-R* | IPSS-M* | Karyotype* | Mutations* | Therapy* | Used for assay** |
|---|---|---|---|---|---|---|---|---|---|
| **MDS1** | female | 60 | MDS-LB | low | low | 46,XX [20] | TET (13%), STAG2 (11%) | none | 1 |
| **MDS2** | female | 75 | MDS-LB | int | moderate high | 46,XX [20] | 3x TET2 (35%/37%/37%), ASXL1 (30%), 2x EZH2 (29%/34%) | Deferasirox | 1 |
| **MDS3** | male | 79 | MDS-LB | int | moderate low | 46,XY [20] | U2AF1 (29%), DNMT3A (8%) | none | 1 |
| **MDS4** | female | 70 | MDS-SF3B1 | very low | very low | 46,XX [20] | SF3B1 (40%) | none | 1 |
| **MDS5** | male | 41 | MDS-SF3B1 | low | low | 46,XY [20] | SF3B1 (43%), NRAS (18%), TET2 (10%) | none | 1 |
| **MDS6** | male | 76 | MDS-SF3B1 | low | low | 46,XY [26] | SF3B1 (37%), DNMT3A (35%) | ESA | 2 |
| **MDS7** | female | 65 | MDS-LB | very low | low | 46,XX [20] | DNMT3A (5%) | none | 2 |
| **MDS8** | female | 84 | MDS-LB | low | high | 46,XX [20] | IDH2 (38%), ASXL1 (16%), SRSF2 (13%), PHF6 (3%) | none | 2 |
| **MDS9** | male | 74 | MDS-LB | low | moderate low | 46,XY [20] | ASXL1 (23%), 2x IDH1 (22%/8%), SRSF2 (21%), NF1 (9%), NRAS (5%) | Prednisolone | 2 |
| **MDS10** | female | 46 | MDS-SF3B1 | low | very low | 46,XX [20] | SF3B1 (38%), CUX1 (2%) | none | 2 |
| **HY1** | female | 60 | hematologically healthy | - | - | - | - | - | 1 |
| **HY2** | male | 53 | hematologically healthy | - | - | - | - | - | 1 |
| **HY3** | female | 63 | hematologically healthy | - | - | - | - | - | 1 |
| **HY4** | female | 84 | hematologically healthy | - | - | - | - | - | 1 |
| **HY5** | male | 76 | hematologically healthy | - | - | - | - | - | 1 |
| **HY6** | female | 87 | hematologically healthy | - | - | - | - | - | 2 |
| **HY7** | male | 58 | hematologically healthy | - | - | - | - | - | 2 |
| **HY8** | male | 53 | hematologically healthy | - | - | - | - | - | 2 |
| **HY9** | male | 43 | hematologically healthy | - | - | - | - | - | 2 |
| **HY10** | female | 65 | hematologically healthy | - | - | - | - | - | 2 |

**Supplementary Figure S1**

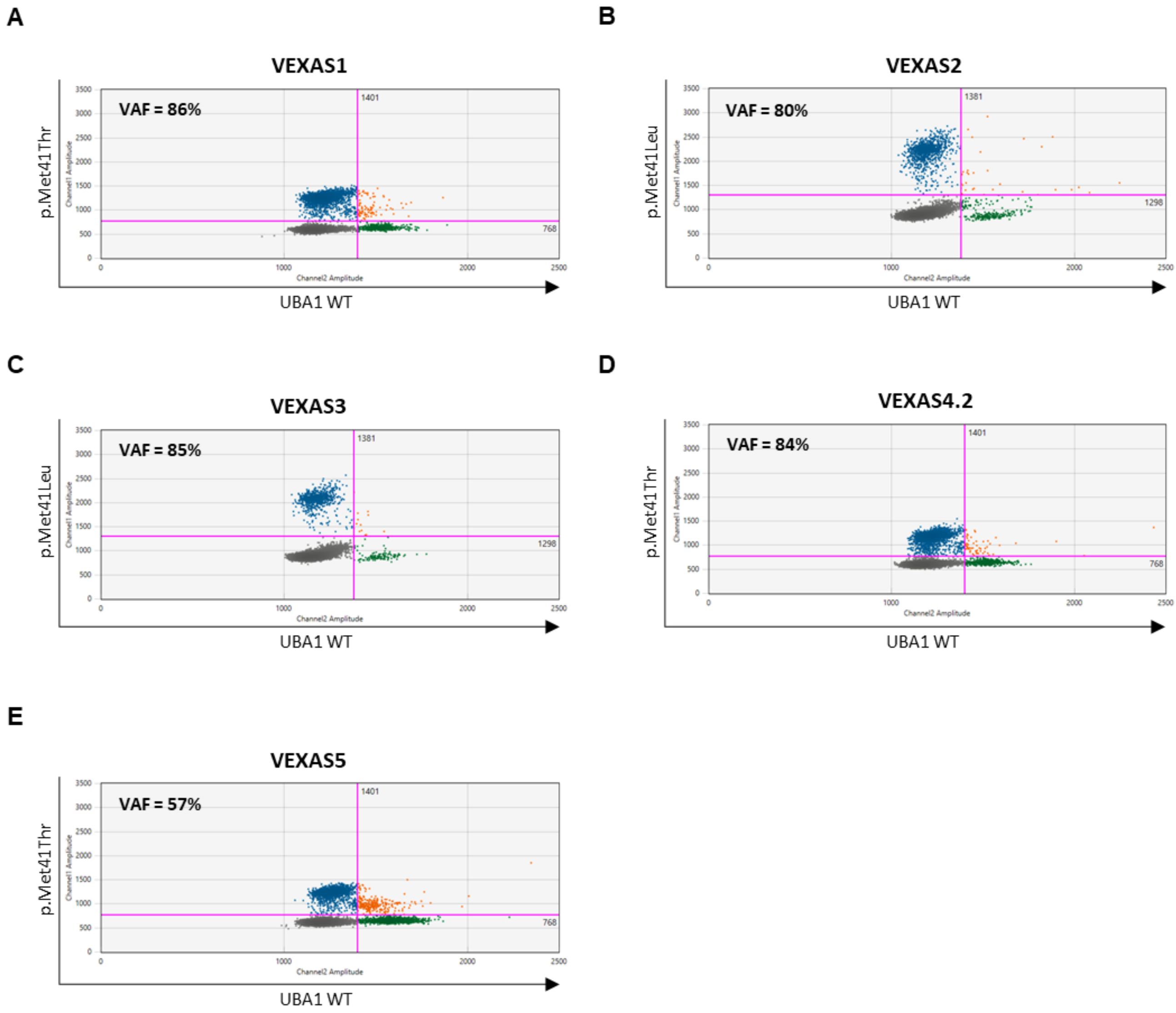


**Supplementary Figure S1. Verification of UBA1 mutations in CD34⁺ cells of VEXAS patients using ddPCR.** DNA from primary CD34⁺ cells of VEXAS patients VEXAS1 (**A**), VEXAS2 (**B**), VEXAS3 (**C**), VEXAS4.2 (**D**) and VEXAS5 (**E**) was analyzed for their previously known *UBA1* mutations using probes for *UBA1 p.Met41Thr* or *UBA1 p.Met41Leu* (y-axis) and UBA1 WT (x-axis). VAF, variant allele frequency.

**Supplementary Figure S2**

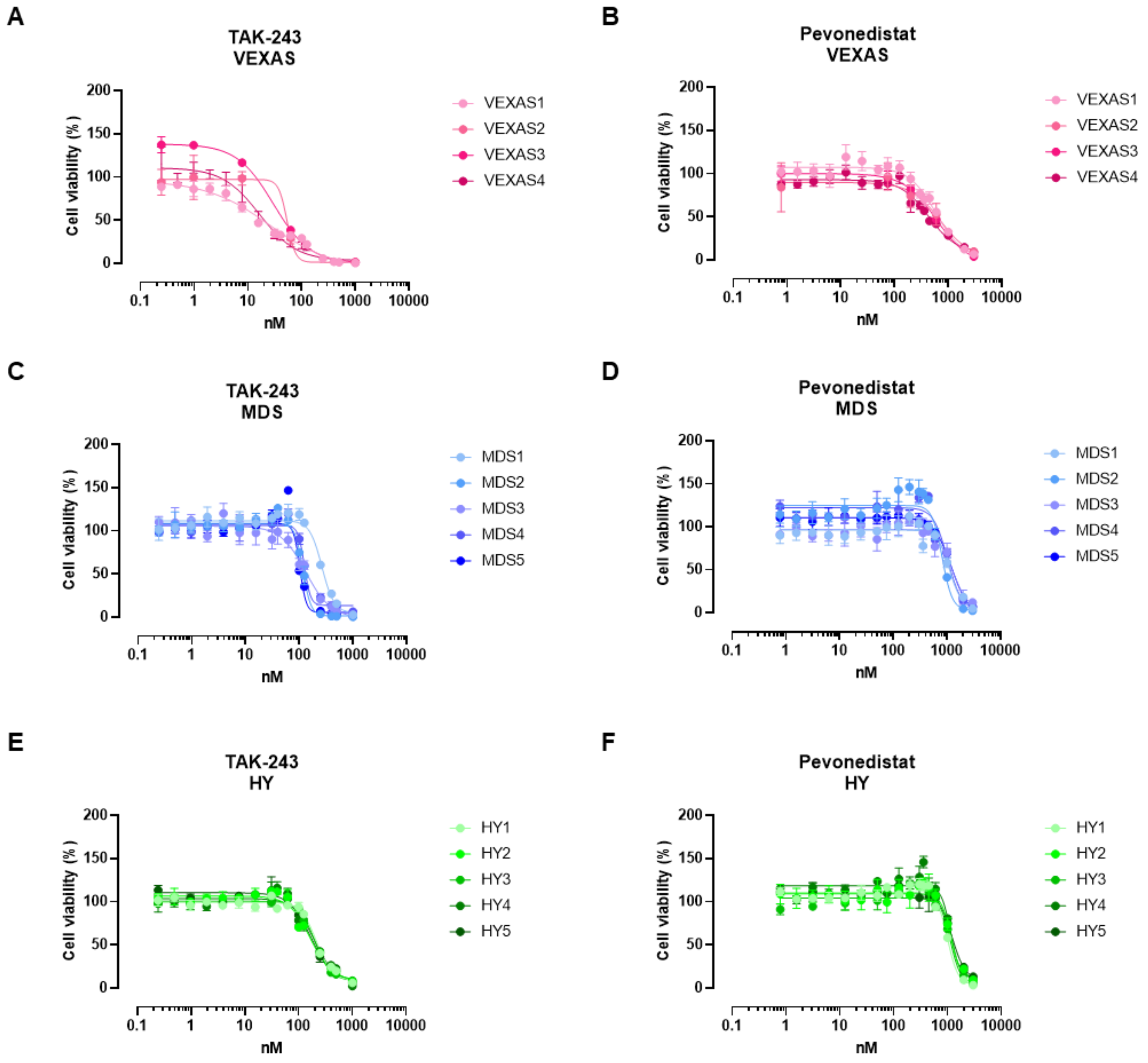


**Supplementary Figure S2. Treatment of VEXAS, MDS and healthy CD34⁺ cells with TAK-243 and pevonedistat.** Dose-response experiments of primary VEXAS (n=4; magenta), MDS (n=5; blue) and healthy (n=5; green) CD34⁺ cells treated with TAK-243 or pevonedistat. Cell viability was measured 48 hours after treatment using CellTiter-Glo® Luminescent Cell Viability Assay. **A**,**C**,**E** Dose response curves of TAK-243 for each VEXAS (**A**), MDS (**C**) and healthy (**E**) CD34⁺ sample. **B**,**D**,**F** Dose response curves of pevonedistat for each VEXAS (**B**), MDS (**D**) and healthy (**F**) CD34⁺ sample. Data were analyzed using nonlinear regression and are represented as mean ± SD.

**Supplementary Figure S3**

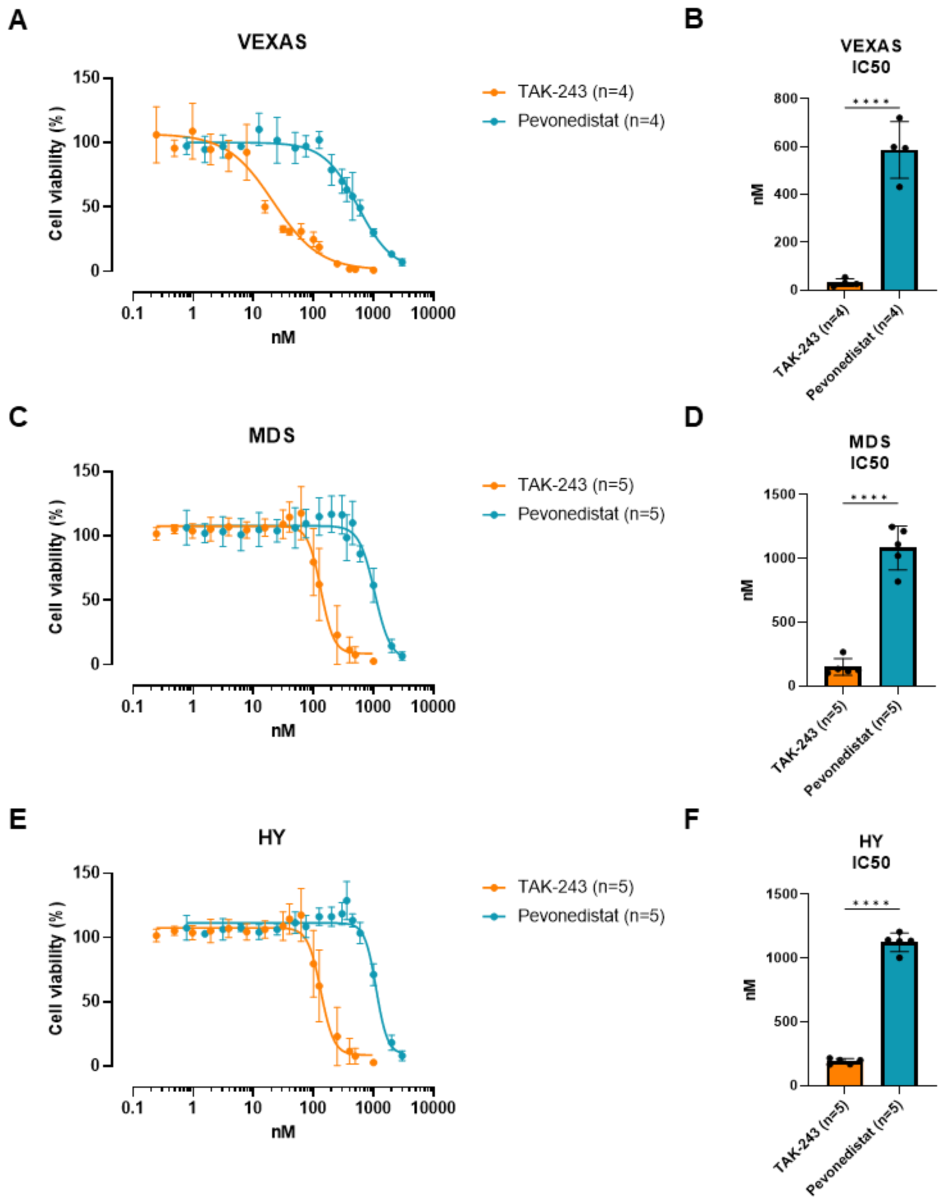


**Supplementary Figure S3. VEXAS, MDS and healthy CD34⁺ cells are more sensitive to treatment with TAK-243 than with pevonedistat.** Dose-response experiments of primary VEXAS (n=4), MDS (n=5) and healthy (n=5) CD34⁺ cells treated with TAK-243 (orange) or pevonedistat (blue-green). Cell viability was measured 48 hours after treatment using CellTiter-Glo® Luminescent Cell Viability Assay. **A**,**C**,**E** Dose response curves of VEXAS (**A**), MDS (**C**) and healthy (**E**) CD34⁺ cells. **B**,**D**,**E** IC50s of VEXAS (**B**), MDS (**D**) and healthy (**F**) CD34⁺ cells. Data were analyzed using nonlinear regression (**A**,**C**,**E**) and two-sided unpaired t-test (**B**,**D**,**F**) using Prism 9 (GraphPad Software), and are represented as mean ± SD. ****$p \leq 0.0001$
